# Comparative analyses of toxin-associated gene homologs from an Old World viper, *Daboia russelii*

**DOI:** 10.1101/152082

**Authors:** Neeraja M. Krishnan, Binay Panda

## Abstract

Availability of snake genome sequences has opened up exciting areas of research on comparative genomics and gene diversity. One of the challenges in studying snake genomes is the acquisition of biological material from live animals, especially from the venomous ones. Additionally, in certain countries, Government permission is required to handle live snakes, making the process cumbersome and time-consuming. Here, we report comparative sequence analyses of toxin gene homologs from Russells viper (Daboia russelii) using whole-genome sequencing data obtained from the shed skin. When compared with the major venom proteins in Russells viper studied previously, we found 45-100% sequence similarity between the venom proteins and their skin homologs. Additionally, comparative analyses of 20 toxin gene family homologs provided evidence of unique sequence motifs in nerve growth factor (NGF), platelet derived growth factor (PDGF), Kunitz/Bovine pancreatic trypsin inhibitor (Kunitz BPTI), cysteine-rich secretory proteins, antigen 5, and pathogenesis-related 1 proteins (CAP) and cysteine-rich secretory protein (CRISP). We identified V11 and T35 in the NGF domain; F23 and A29 in the PDGF domain; N69, K2 and A5 in the CAP domain; and Q17 in the CRISP domain to be responsible for differences in the largest pockets across the protein domain structures in crotalines, viperines and elapids from the in silico structure-based analysis. Similarly, residues F10, Y11 and E20 appear to play an important role in the protein structures across the kunitz protein domain of viperids and elapids. Our study sheds light on the usefulness of studying venom protein homologs from skin, their unique features and evolution in vipers. Data deposition: Russells viper sequence data is deposited in the NCBI SRA database under the accession number SRR5506741 and sequences for the individual venom-associated gene homologs to GenBank (accession numbers in Table S1).

## I. INTRODUCTION

Snake venom genes and their products offer an excellent model system to study gene duplication, evolution of regulatory DNA sequences, and biochemical diversity and novelty of venom proteins. Additionally, snake venoms have tremendous potential in the development of new drugs and bioactive compounds (Vonk et al. 2011). Previous studies have highlighted the importance of gene duplications and/or sub-functionalization (Hargreaves et al. 2014; Malhotra et al. 2015; Rokyta et al. 2011) and transcriptional/post-transcriptional mechanisms (Casewell et al. 2014) contributing towards snake venom diversity. Venom studies, so far, have extensively used data from proteomics experiments alongside individual gene sequences or sequences of particular family members to study variations on gene structure and sequence composition. Presently, whole genome sequences of several snake species, king cobra *Ophiophagus hannah* (Vonk et al. 2013); Burmese python *Python bivitattus* (Castoe et al. 2013); rattlesnake *Crotalus atrox* (Dowell et al. 2016); Florida pygmy rattlesnake *Sistrurus miliarius barbouri* (Vicoso et al. 2013); garter snake *Thamnophis elegans* (Vicoso et al. 2013); five-pacer viper *Deinagkistrodon acutus* (Yin et al. 2016); *Protobothops mucrosquamatus* (NCBI Accession PRJDB4386); and corn snake *Pantherophis guttatus* (Ullate-Agote et al. 2014), have either been published or their sequence are made available in the public domain. In addition, genome-sequencing efforts are either underway or the sequences of venom-associated genes have been deposited in the databases for a few others (Kerkkamp et al. 2016). Out of the sequenced genomes, only a few have been annotated, or the annotations have been made public, a key requirement for comparative analysis of genes. This, along with the lack of availability of whole genome sequences and/or complete transcript sequences from venom glands for most snakes has limited studies on toxin gene orthologies and gene variation among venomous snakes.

Four snakes, Russell’s viper (*Daboia russelii*), saw-scaled viper (*Echis carinatus*), spectacled cobra (*Naja naja*), and common krait (*Bungarus caeruleus*) are responsible for most snakebite-related mortality in India (Mohapatra et al. 2011; Warrell et al. 2013; Whitaker 2015). Russells viper is a member of the taxon Viperidae and subfamily Viperinae and is responsible for large numbers of snakebite incidents and deaths in India. Very little is known about the diversity of genes from any viper, including the only viperine where complete genome sequence information is available (European adder, *Vipera berus berus*, https://www.ncbi.nlm.nih.gov/bioproject/170536). Lack of complete genome annotation from this viper using transcripts obtained from venom glands and other snake species reduces the scope of a detailed comparative study on genes, including the toxin-associated genes. Such a study involving various groups of venomous and non-venomous snakes, in addition to other venomous vertebrates and invertebrates, will facilitate our understanding on the evolution of these genes, their diversity, and function.

One of the challenges in studying the genomes of venomous animals is related to sample acquisition. Additionally, in India, Government permission is required to catch snakes and extract blood samples from them (all snakes are protected in India under the Indian Wildlife Protection Act, 1972). This may be circumvented by the use of skin exuviate (shed skin) that does not require drawing blood or taking any tissue from the animals. However, working with DNA isolated from shed skin has its own challenges. Microbial contamination, lack of full-length DNA in the exuviate cells, rapid degradation of DNA in humid conditions and computational challenges in dealing with short stretches of DNA are some of the bottlenecks for working with DNA from exuviate skins.

In the current study, we explored the possibility of getting toxin gene homolog information from low-coverage whole-genome sequencing data using shed skin from Russells viper, and performed comparative analysis on the major toxin proteins from a previously studied report with their predicted skin homologs representing all the 20 venom-associated protein families (Fry 2005). We used the coding sequences and annotation from a previously characterized crotalin, a pit viper, *Protobothrops mucrosquamatus* for the analysis. On the venom homologs, we focused our analyses on five key protein domains; nerve growth factor (NGF), platelet derived growth factor (PDGF), Kunitz/Bovine pancreatic trypsin inhibitor (Kunitz BPTI), cysteine-rich secretory proteins, antigen 5, and pathogenesis-related 1 proteins (CAP) and cysteine-rich secretory protein (CRISP) in Russells viper. Our study identified venom homologs from skin that are highly similar to the venom proteins and the key residues that are changed across the members of viperinae, crotalinae and elapidae that might have contributed towards the evolution of venom in vipers.

## II. MATERIALS AND METHODS

### A. Russells viper shed skin and DNA isolation

Freshly shed skin of Russells viper from Bangalore, India was a gift from Mr. Gerry Martin. The skin exuviate for the entire snake was obtained, cleaned thoroughly with 70% ethanol and with nuclease-free water 3 times each, dried thoroughly and frozen until the time of extraction of DNA. Genomic DNA was extracted following the protocol of Fetzner (Fetzner 1999) with modifications.

### B. Sequencing, read processing and assembly

Illumina paired-end read libraries (100bp paired-end reads with insert size of 350bp) were prepared following the manufacturer instructions using amplification free genomic DNA library preparation kit and sequenced using Illumina HiSeq2500 instrument. Archaeal, bacterial and human sequence contamination were removed from the Russells viper sequence by DeConSeq (Schmieder & Edwards 2011) using curated and representative genomes (https://www.ncbi.nlm.nih.gov/genome/browse/reference/).

Furthermore, the sequenced reads were post-processed to remove unpaired reads and quality analysis was performed using FastQC v0.1 (http://www.bioinformatics.babraham.ac.uk/projects/fastqc/). The rd len cutoff option was exercised during the read assembly step to trim off the low-quality bases, since the per-base quality was found to drop below 28 after the initial 50-70 bases of the read. The Russells viper read libraries were assembled using SOAPdenovo2 (r240) (Luo et al. 2012).

### C. Identifying toxin gene homologs, coding regions, and predicted gene structures

The DNA sequences for 51 out of 54 venom-associated genes (Fry 2005) from Protobothrops mucrosquamatus were downloaded (Table 1). These were used to fish genomic scaffolds bearing highly similar sequences in Russells viper genome assembly, using BLAST with an E-value threshold of 103. The fished scaffolds were then anchored to the respective coding sequences from Protobothrops mucrosquamatus using a discontiguous megaBlast, to determine the correct frame of translation and extract the complete amino acid coding sequence (CDS) corresponding to Russells viper toxin-gene homologs. We obtained the exon-intron structures for all the toxin-associated gene homologs in Russells viper by aligning the CDS with gene sequences using discontiguous megaBlast and plotted using the tool GSDS2.0 (Hu et al. 2015). The sequences for the Russells viper venom-associated gene homologs were deposited in GenBank and their accession numbers for are provided in Table S1.

**Table 1:**
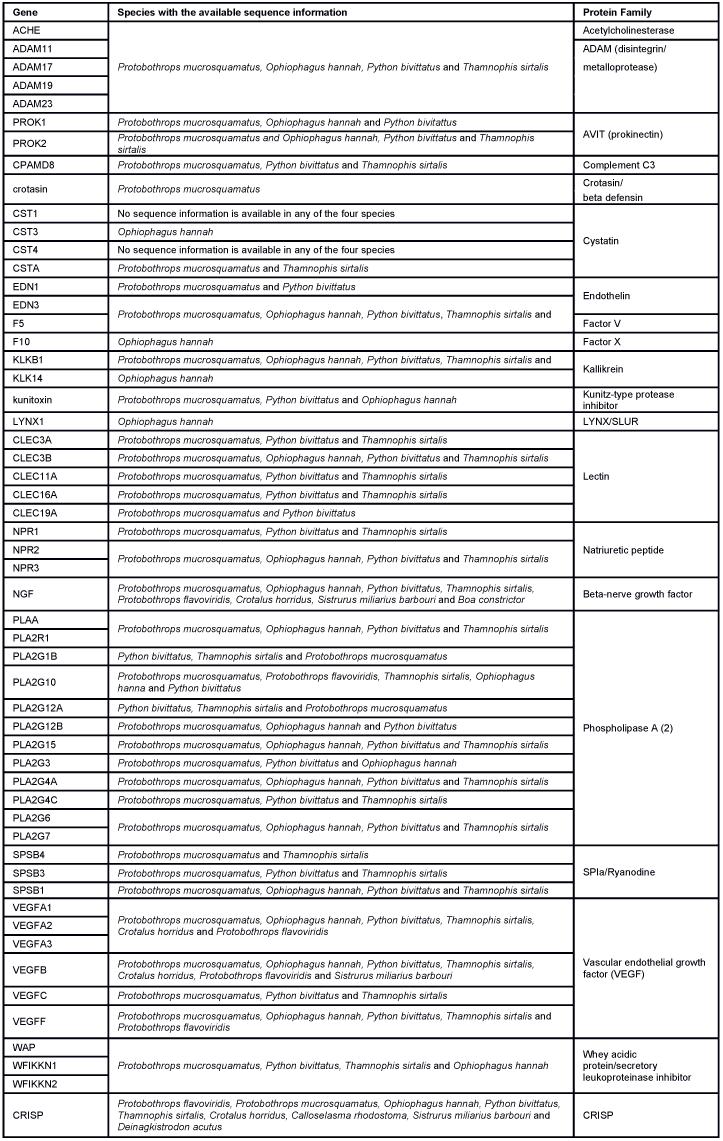
Genes and their representative families used in the current study.

### D. Comparative analysis between venom proteins and their skin homologs

We obtained the accession IDs for the major toxin families from Russells viper of Indian sub-continent (Supplementary Figure S1 in (Sharma et al. 2015). Their corresponding protein sequences were matched using blastp with the amino acid sequences from the derived skin homologs. For the genes covered under each family, a percent identity metric, indicative of the extent of sequence similarity between the venom proteins and their skin homologs, was estimated. Similar comparative analyses were performed for king cobra (Ophiophagus hannah) using accession IDs provided in Additional File 4 of Tan et al. 2015, and the predicted toxin-associated protein homologs from blood of king cobra (PRJNA201683; (Vonk et al. 2013). Comparative analyses were performed using blastp, with the venom protein sequence as the query, against PRJNA201683.

### E. Comparative analyses of venom protein homolog domains

The amino acid sequences of all the Russells vipers toxin-associated gene homologs were subjected to domain search using Pfam (Finn et al. 2016) (Table S2). All domain sequences were aligned using blastp to non-redundant protein sequences from 18 snake species (Table S3). We wanted to compare the gene structures of venom-associated gene homologs between the venomous and the non-venomous animals, hence included sequence information from members of the later group. Five domains (NGF, PDGF, Kunitz BPTI, CAP and CRISP) from four genes (NGF, VEGF, CRISP/Serotriflin, and Kunitoxin), with variability across different snake groups and where sequence information were available beyond the whole genome sequences, were used for expansive comparative analyses (Table S4) using sequences from Viperids (taxid: 8689), elapids (taxid: 8602), Colubrids (taxid: 8578), Boids (taxid: 8572), Acrochordids (taxid: 42164), Pythonids (taxid: 34894), lizards (squamates (taxid: 8509) minus snakes (taxid: 8570)), Crocodylia (taxid: 51964) and Testudines (taxid: 8459).

### F. 3D structure prediction of the chosen domains

Consensus sequences were determined from NGF, PDGF, Kunitz BPTI, CAP and CRISP domain alignments using Simple Consensus Maker (https://www.hiv.lanl.gov/content/sequence/CONSENSUS/SimpCon.html) for crotalines (CR), viperines (VP) and elapids. The consensus sequences were submitted to the protein fold recognition server (Kelley et al. 2015) using standard mode (http://www.sbg.bio.ic.ac.uk/phyre2/html/page.cgi?id=index). The best 3D model was further investigated by Phyre2 to analyze the structural model using various open source tools.

## III. RESULTS

### A. Shed skin yielded fairly good quality DNA for genome sequencing and near complete coding sequences for toxin-associated gene homologs

Genomic DNA isolated from the shed skin of Russells viper was fairly intact with most of the DNA in the size range of more than 5kbp (Fig. S1). Sequenced short reads were assembled and then used to fish all the 51 toxin-associated genes in Russells viper (see Materials and Methods). Next, we obtained the exon-intron structures for all homologs in Russells viper by aligning the CDS with gene sequences (Fig. S2). We found the average length of the exons in Russells viper toxin-associated gene homologs to be around 190 nucleotides (nt), matching well with the lengths of other vertebrate exons (Gelfman et al. 2012).

### B. Similarity between venom proteins and their predicted skin homologs

For the Russells viper, we found 45 - 100% sequence similarity between the major venom proteins and their predicted skin homologs (Figure 1). The sequences for venom nerve growth factor (VNGF) and its skin homolog were identical. Similarly, VEGF and CRISPs from venom gland were highly similar to their skin homologs (99% and 92% sequence similarity respectively). Other proteins like, KSPI, SVSPs and PLA2 showed 79%, 74% and 61% sequence identity, respectively (Figure 1 and Fig. S3). In order to find out whether the sequence divergence between some of the venom gland proteins and their predicted skin homologs was specific to Russells viper, we performed similar analysis using venom proteins and their blood homologs from king cobra, Ophiophagus hannah (Vonk et al., 2013). In the case of Ophiophagus hannah, the differences between toxin proteins and their blood homologs were minor for most families studied (similarity ¿= 75%), except for PLA2, which had a low similarity of 23% (Fig. S4 and Fig. S5).

**Figure 1:**
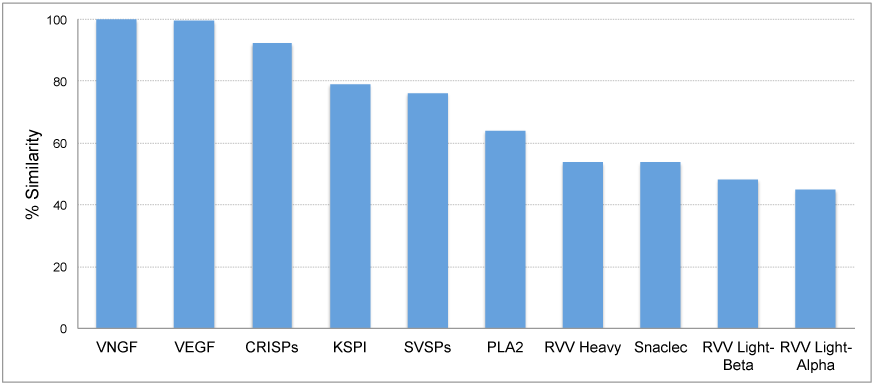
Sequence identity (%) between the proteins from ten major venom families and their skin homologs in Russells viper. The homolog with the highest identity was considered in cases with more than one homolog.

### C. Comparative domain analyses of venom protein homologs

Among the genes, a larger pool of sequences were available only for NGF, PDGF domain of VEGF, Kunitz BPTI domain of Kunitoxin, CRISP and CAP domains in CRISP and Serotri-flin proteins, from various snake groups (Colubridae, Boidae, Pythonidae and Acrochordidae), non-snake reptilian groups (lizards, crocodiles and Testudines), venomous invertebrates (wasps, spiders and scorpions) and venomous vertebrates (fishes and mammals). Therefore, these domains were compared with those from Russells viper. Comparative domain analysis was performed for all toxin-associated gene homologs (Fig. S6) across 18 snake species where sequence information was available (Table S3). In the case of five domains: CAP and CRISP domains of CRISP and serotriflin genes (L and AL), Kunitz BPTI of kunitoxin (S), NGF (T) and PDGF of VEGFA (AP-AR) and VEGFF (AU), we found that the maximum number of species aligned to their domain sequences. Some protein domains, the CRISP, Kunitz BPTI, guany-late CYC, PDGF of VEGFF and WAP, showed long stretches of mismatches (Fig. S6) compared with Russells viper sequence. Out of these, only NGF and PDGF domains of VEGF had amino acid changes specific to the members of the group crotalinae, that were completely absent in any other group used for comparison, including in lizards, crocodiles, and turtles (Fig. S7). Specific changes in these proteins and their implications are discussed below.

Russells viper NGF gene homolog is a single exon gene with a 745nt transcript coding for a 244 amino acid protein consisting of a single NGF domain (Figure 2A). The NGF domain bears 28% sequence conservation across all the five vertebrate phyla, namely, fishes, amphibians, reptiles, aves and mammals distributed along the length of the domain (Figure 2B). Thirty-six percent out of these residues are conserved across other venomous vertebrates (fishes, squamates and mammals) and venomous invertebrates (scorpions and wasps) (Figure 2C). Thirteen percent of Russells viper NGF domain residues are variable with respect to the domain sequence in at least one among the NGF sequences in the groups of vipers and elapids (Figure 2D). Although several amino acids in the NGF domain in New World vipers seem to have changed from the Russells viper and other vipers of the group viperinae, their function probably remains unchanged. For example, phenylalanine (F) to isoleucine (I) at position 12 and serine (S) to asparagine (N) at position 19 between the crotalines and viperines does not change the function of the amino acids (from one hydrophobic amino acid to another and from one polar amino acid to another). However, there are others, for example, threonine (T) and glutamine (Q), at position 67 and 68 respectively in the NGF domain of the New World viper, which were only there in that specific group. One of those, a polar amino acid glutamine at position 68, is a very important residue as its corresponding amino acid in any of the other snakes, except in colubrids, is a hydrophobic proline.

**Figure 2:**
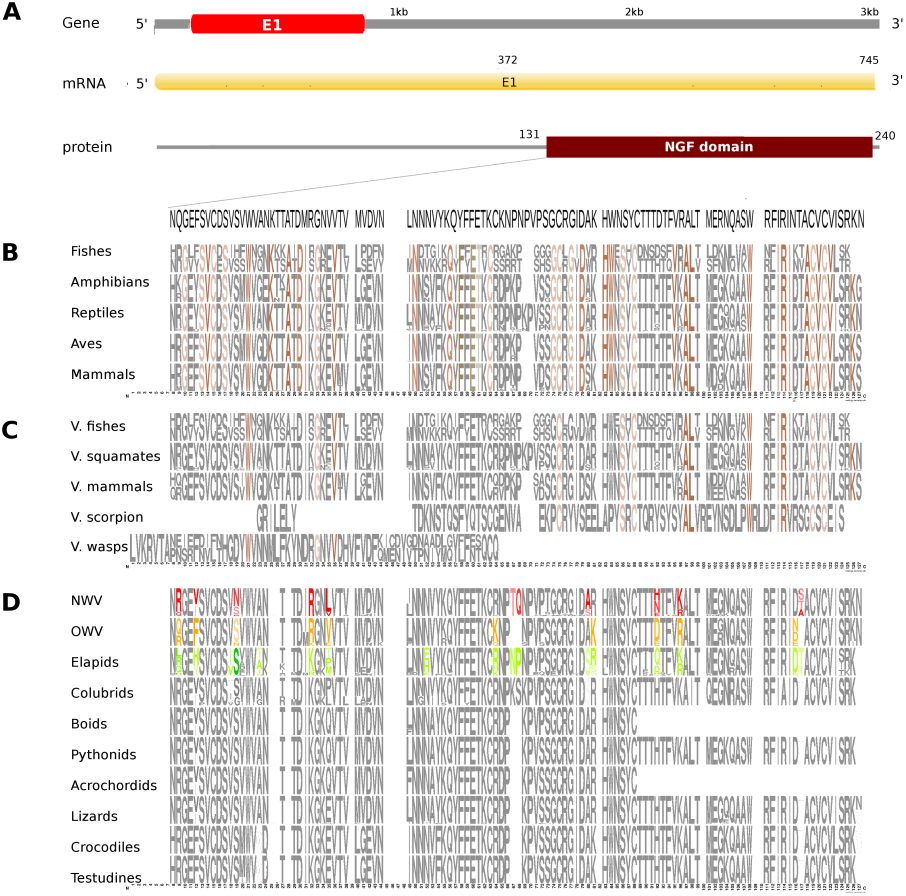
Comparative analyses of nerve growth factor (NGF). NGF gene, mRNA, and protein domains of Russell’s viper (A) and its comparison with the consensus NGF sequences from all five vertebrate phyla (fishes, amphibians, reptiles, birds and mammals) (B), with venomous (V) vertebrates from multiple phyla of vertebrates and invertebrates (C), and from various reptilian subgroups (D) are shown. The shades of brown and grey in B and C represent conservation to various degrees and variability, respectively. Grey in D represents conserved residues, red represents variable residues in the crotalines (CR), yellow and green represent conserved and variable residues in the viperines (VP), and elapids respectively.

In Russells viper, the VEGFA gene homolog comprises five exons coding for a 652nt long transcript and a protein with two domains: PDGF and VEGF-C (Figure 3A). The PDGF domain sequence exhibits conservation in 65% of its residues across the three vertebrate phyla (reptiles, birds and mammals) (Figure 3B). Since sequence information from fishes and amphibians were not available, they could not be included in the comparison study. Out of the conserved residues, 21% of those were also conserved in venomous vertebrates (squamates and mammals) and venomous invertebrates (wasps). Fifteen percent of the PDGF domain residues were variable in at least one of the two snake groups: vipers and elapids (Figure 3C and Figure 3D). Like the NGF domain, the evolution of the PDGF domain in New World vipers at certain amino acids is striking. For example, in the crotalins, the position 67 is a polar amino acid tyrosine (Y) while in all other reptiles, venomous invertebrates and mammals; this is primarily a hydrophobic amino acid phenylalanine (F). This might bear implications on the proteins structure and function.

**Figure 3:**
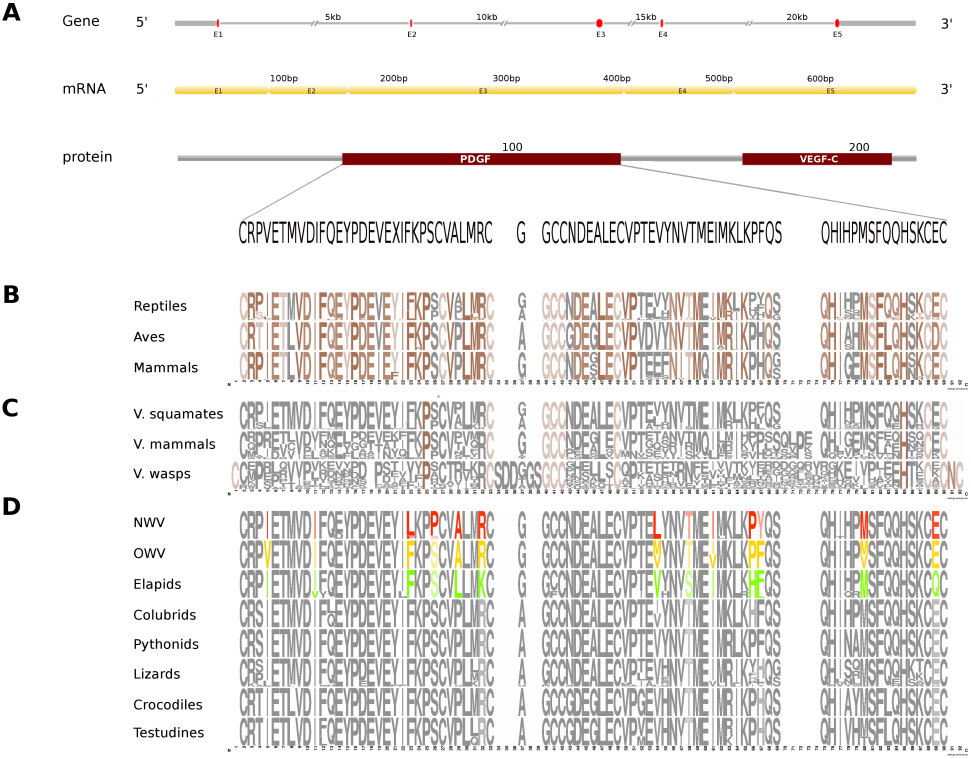
Comparative analyses of vascular endothelial growth factor - A (VEGF-A). Organization of the gene, mRNA, and protein domains of Russells viper PDGF domain (A) and its comparison with the consensus sequences from all five vertebrate phyla (fishes, amphibians, reptiles, birds and mammals) (B), from the venomous (V) vertebrates and invertebrates (C), and from various reptilian subgroups (D) are shown. The shades of brown and grey in B and C represent conserved and varying residues, respectively. Grey in D represents conserved residues, red represents variable residues in the crotalines (CR), yellow and green represent conserved and variable residues in viperines (VP), and elapids respectively.

Kunitoxin gene homolog in Russells viper is a 3.1kb gene comprising two exons, with a transcript length of 270nt that codes for a 44 amino acids long single Kunitz BPTI domain (Figure 4A). About 29% of the protein domain residues are conserved across the four vertebrate phyla (amphibians, reptiles, aves and mammals) (Figure 4B). Since sequence information from the Kunitz BPTI for fishes was not available, they could not be included in the comparison. Out of these conserved residues, 76% are conserved in venomous vertebrates (squamates and mammals) and venomous invertebrates (scorpions and wasps) (Figure 4C) and 56% of the domain residues are variable in at least one of two snake groups (vipers, and elapids) (Figure 4D). Of the residues that are evolved in the members of crotalinae, the second residue, a positively charged one, alanine (A) is present only in the members of viperinae, which is replaced by a hydrophobic residue, proline (P), in the crotalines and elapids. Residues 14-18 are very polymorphic in the crotalines and elapids, but not so in the viperines.

The CRISP gene homolog in Russells viper is a 25kb long gene, comprises of 8 exons coding for a 787nt transcript and two protein domains, CAP and CRISP (Figure 5A). The CAP domain exhibits conservation in 7% of its residues across all the five vertebrate phyla (Figure 5B). Forty-two percent of those residues are conserved across venomous vertebrates (amphibians, squamates and mammals) and venomous invertebrates (scorpions and wasps) (Figure 5C). In addition, there are five residues conserved across all the venomous animals (Figure 5C). Twenty-seven percent of the CAP domain and 15% of the CRISP domain residues are variable in at least one of the three snake groups (Figure 5D). There are several extra residues for the CAP domain in the crotalines and elapids, but not in the viperines. The conserved residues comprised mostly of Cystines and to a lesser extent Asparagines (Figure 5E) across venomous vertebrates (squamates and mammals) (Figure 5F). Sixty percent of the CRISP domain residues are variable in at least one viperine or elapid member with respect to the domain sequence of Russell’s viper (Figure 5G).

**Figure 4:**
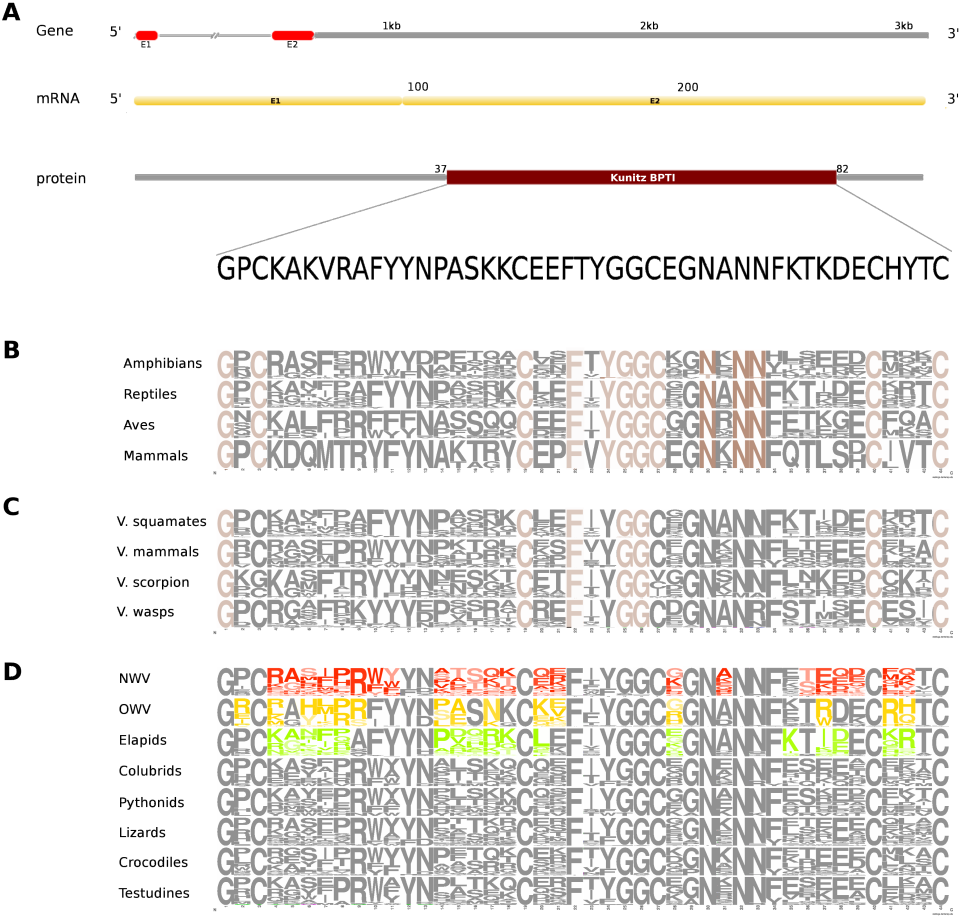
Comparative analyses of kunitoxin. Organization of the gene, mRNA, and protein domains of Russells viper (A) and its comparison with the consensus BPTI domain sequences from all five vertebrate phyla (fishes, amphibians, reptiles, birds and mammals) (B), from venomous (V) vertebrates and invertebrates (C), from various reptilian subgroups (D) are shown. The shades of brown and grey in B and C represent conserved and varying residues, respectively. Grey in D represents conserved residues, red represents variable residues in the crotalines (CR), yellow and green represent conserved and variable residues in viperines (VP), and elapids respectively.

**Figure 5:**
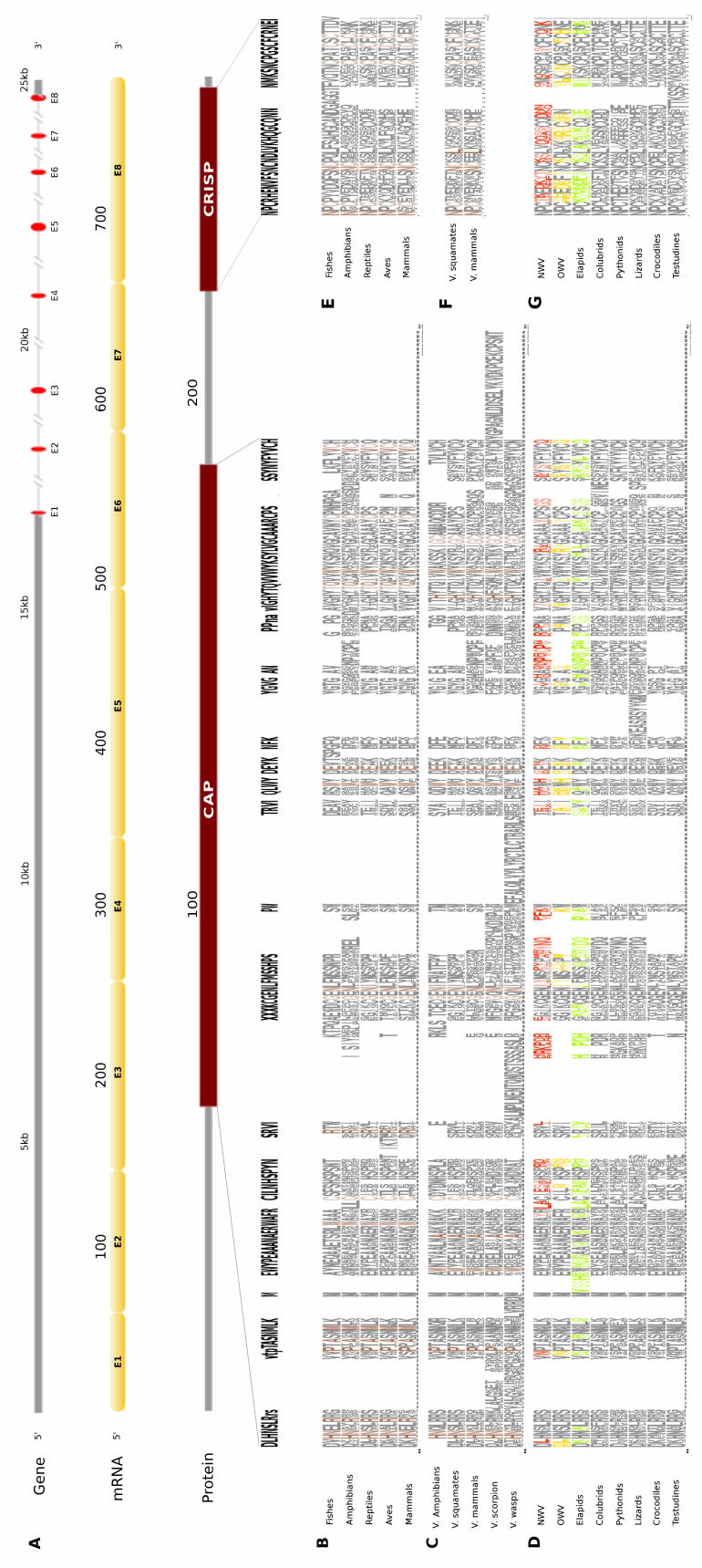
Comparative analyses of CRISP. Organization of CRISP gene, mRNA, and protein domains of Russells viper (A) and its comparison with the consensus CRISP sequences from all five vertebrate phyla (fishes, amphibians, reptiles, birds and mammals, B and E); from venomous animals (V) vertebrates (fishes, squamates and mammals) and invertebrates (scorpions and wasps, C and F); and from various reptilian subgroups (D and G) are shown. The shades of brown and grey in B, C, E and F represent conserved and varying residues, respectively. Grey in D and G represents conserved residues, red represents variable residues in the crotalines (CR), yellow and green represent conserved and variable residues in viperines (VP), and elapids respectively.

Next, we explored the role of consensus domain sequences and their possible role of conserved amino acids in those key toxin-associated protein domains across vipers and elapids. We constructed the 3D structure models using Phyre2, followed by Phyre2 investigation, for further analyses on the structural model. As evident from the analyses, amino acid residues 18-19 and 117 of the NGF domain reflected a difference in mutation sensitivity as detected by SusPect algorithm (Yates et al. 2014), especially in the elapids compared to the viperids (Figure 6). Residue 18 is Valine in the viperines and Isoleucine in the elapids; residue 19 is Serine in the viperines and Asparagine in the crotalines; and residue 117 is Threonine in the elapids and Serine in the crotalines (Figure 6A). This might have implications in the structure of the protein as the largest pockets detected by fpocket algorithm appear to be vastly different among the crotalines, viperines and elapids for the NGF, PDGF, CAP and CRISP domains (Figure 6). The pockets appeared small in all cases for the elapids, and largest in the case of viperines (Figure 6). Minor differences in clashes were observed at residues 10,11 and 20 of the Kunitz domain and residue 38 of this domain showed a rotamer conflict in the case of the New World vipers (Figure 6C). Similarly, residue 46 of the CAP domain and residues 4 and 31 of the CRISP domain showed rotamer conflict for the viperines (Figure 6D and Figure 6E). The other protein quality and functional parameters were not affected across the 3D structure models for the three snake groups (Fig. S8).

**Figure 6:**
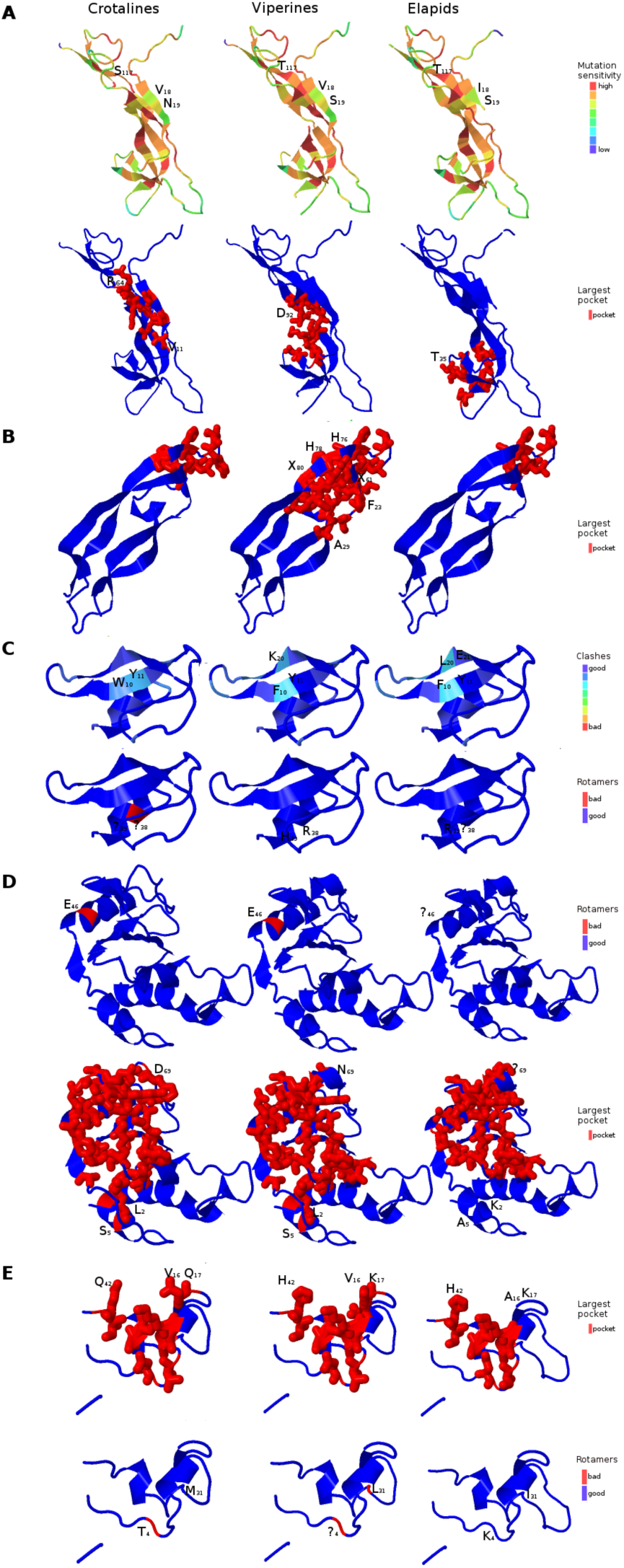
Three-dimensional protein structural models to access quality and functional differences in key venom-associated proteins. NGF, A; PDGF, B; Kunitz BPTI, C; CAP, D; and CRISP, E) across crotalines (CR), viperines (VP) and elapids are shown. The status of the parameters being investigated using Phyre2 are indicated in the color legends on the side.

## IV. DISCUSSION

Accessibility and affordability of high throughput sequencing technologies along with the availability of sophisticated computational tools to assemble, annotate and interpret genomes is playing a powerful role in deciphering gene functions and their role in evolution. Snake toxin genes are coded by gene families and produce gene isoforms through the process of duplications (Casewell et al. 2013; Fry 2005). Several studies on the venom-associated proteins from New World vipers have classified the venoms into four groups (type I-IV), based on the relative abundance of toxin families (Calvete 2013; Gibbs et al. 2013; Goncalves-Machado et al. 2016; Jimenez-Charris et al. 2015; Lomonte et al. 2014; Mora-Obando et al. 2014; Pla et al. 2017; Salazar-Valenzuela et al. 2014). The different groups are: snake venom metalloproteinase-predominant (type I), heterodimeric-neurotoxic PLA2 rich (type II), serine proteinases and PLA2 (type III) and type IV, which is similar to type III but with significant higher concentration of snake venom metalloproteinases (Calvete 2017). Russells viper (Daboai russelli) is a Old World pitless viper, characterized by the lack of heat sensing pit organs (Mallow et al. 2003). There is significant variation in the venom composition of Russells viper in India (Jayanthi & Gowda 1988; Sharma et al. 2015) making anti-venom produced using snake venoms from a single location against all Russells viper bites across the country ineffective. The variation in the venom composition within the same species is thought to be a result of adaptation in response to the difference in diets (Barlow et al. 2009; Casewell et al. 2013; Daltry et al. 1996). Currently, efforts are underway to collect venoms of Russells viper from different regions of India in order to understand the differences in their venom composition (Rom Whitaker, and Gerry Martin, personal communications).

Studies on venom-associated genes or their homologs using whole-genome sequencing data in Russells viper are scarce. Past studies on the members of viperadae focused on proteins and used proteomics-based analyses (Gao et al. 2013; Gao et al. 2014; Kalita et al. 2017; Li et al. 2004; Liu et al. 2011; Mukherjee et al. 2016; Sharma et al. 2015; Tan et al. 2017; Tan et al. 2015; Villalta et al. 2012). Currently, information on gene homologs from any viper, especially Russells viper, is limited and therefore, comparative analyses between the toxin genes and their homologs will add value to our understanding. The only viperine where complete genome sequence information is available is a European adder, Vipera berus berus (https://www.ncbi.nlm.nih.gov/bioproject/170536). Although sequence information is available for this species, the annotation is not available and therefore could not be used in our study.

The aim of the current study was two fold. First, as handling and getting biological material from snakes in India requires Government permission and specific expertise, we wanted to test whether good quality whole-genome sequence information can be obtained using shed skin. Second, we wanted to see which toxin-associated gene homologs are true surrogates of their venom gland-derived proteins through comparative genomics study. On the first account, we found the results to be satisfactory. Although shed skin is often contaminated with bacteria and other microorganisms and the DNA obtained from the shed skin is sheared, using freshly shed skin, we successfully isolated high molecular weight genomic DNA (Fig. S1), which was subsequently used to generate genome sequencing data. We believe that shed skin may be an attractive option for generating snake genome data and studying molecular evolution. On the second account, and in order to demonstrate that skin-derived toxin-associated protein homologs can add value to toxinology studies, we compared previously studied Russells viper venom proteins (Sharma et al. 2015) with their skin-derived predicted venom-associated homologs. Results from our analyses showed that some of the venom gland proteins are identical or near identical to their skin-derived homologs (VNGF, 3FTX and LAAO) but others had low overall similarity (Snaclec and RVV). We were curious to find out whether the low sequence similarity for some venom proteins with their homologs was specific to Russells viper and how much of the low overall similarity in those proteins was due to the heterogeneity, if any, found among snakes of the same species. Comparative analysis between the toxins and their blood homologs in king cobra provided us with an answer for the first question where 7 out of 8 venom proteins studied (except for PLA2) were very similar to their blood homologs (Fig. S4 and Fig. S5). This suggests that some venom proteins may not be that different from their homologs in other organs. Strength to this hypothesis comes from a recent study in python where the authors argue that not the expression but the functional evidence of toxic effects on prey is the correct criterion to classify proteins as venom toxins (Reyes-Velasco et al. 2015). However, we are aware that this may vary from species to species and in some species the venom proteins may be very different from their homologs. Our data suggests that the degree of variation between the venom toxins and their homologs is greater in Russells viper than in king cobra. When we compared the 4 published studies (Kalita et al. 2017; Mukherjee et al. 2016; Sharma et al. 2015; Tan et al. 2015) on Russell’s viper venom proteins, we found that the composition of some of the major venom proteins varied significantly (Figure S9). For example, in one study (Mukherjee et al. 2016), VNGF constituted only 0.4% of the venom while in another (Kalita et al. 2017), the same protein constituted 4.8% of the venom. As both studies came from the same lab, there is little chance for any technical or assay-related variability. In the first study, the venom was used from the captive species in a zoo in the USA from Pakistani origin (Mukherjee et al. 2016) while the other used venom from a commercial source in India (Kalita et al. 2017). This suggests that there is a great deal of variation in the composition of Russells viper venom collected from different locations, corroborating the earlier results from a different group (Jayanthi & Gowda 1988; Sharma et al. 2015). In our study, we compared venom proteins described previously (Sharma et al. 2015) using animals captured near Chennai, Tamil Nadu, India with skin-derived homologs from a completely different animal (shed skin was collected in Bangalore, Karnataka, India). The distance between these two places is roughly 350-400km. Therefore, it is possible that in our study, the low similarity in some of the venom proteins with their skin homologs could have been due to the variation in the venom in the animals in these two locations. Despite this, 50% of the venom proteins studied had ¿75% and 3 had near perfect sequence similarity with their skin homologs. A clear picture will emerge from a direct comparison between the venom proteins and their skin homologs from the same animal.

From the sequence data, we succeeded in assembling near complete CDS for 20 gene families representing 51 gene homologs (Fry 2005). This highlights the utility of genome sequencing data in studying toxin gene homologs. As the lengths of the toxin-associated gene homologs in Russells viper were much longer than the CDS, the intronic sequences were assembled with gaps. This was primarily due to the low coverage sequencing data used for assembly and the lack of long-insert mate pair sequencing data in our repertoire. This was not a problem as the aim of our study was to study toxin protein homologs and not to assemble Russells viper genome. The mean length of exons for the toxin-associated genes in Russells viper was 190 base pairs, much smaller compared to the average intron length. In our study, we could assemble exons accurately using short-read sequence data. Interestingly, we found that the AT to GC ratio in the CDS regions (cumulatively for all the 51 genes) of the toxin-associated gene homologs was 1:1 whereas it was skewed (the ratio is 1.5:1) for the full gene sequences.

Despite the advantages of our study, it has certain limitations. First, we studied toxin gene homologs and not the toxin genes. Therefore, there is a possibility that the toxin genes from the same animal are different from their homologs in the skin. Studying skin and venom gland-derived DNA and protein and from the same animal alongside the functional studies shall provide a definitive conclusion in this regard. Second, like any other annotation-based study, our study relies on the quality of existing/prior annotation of toxin-related genes. Although the chances are slim, as we used sequence information from a crotaline to extract the toxin-associated gene sequences in Russells viper, it is entirely possible that we might have missed some key genes specific to Russells viper. Finally, we did not study the biological relevance of specific mutations in the toxin gene homologs in the viperine. Future studies using high-coverage sequencing data to derive better gene annotation along with the studies on their spatial, temporal expression will point to the true functional significance of toxin gene homologs.

## V. CONCLUSIONS

Our results demonstrated the feasibility of obtaining good quality genomic DNA using freshly shed skin from venomous snakes to generate *de novo* sequencing data. This will aid sequencing, analyses and interpretation of their genomes without catching the animals. Additionally, our study demonstrated that it is possible to obtain information on some venom toxins using their predicted gene homologs from other organs.

